# The measure of spatial position within groups that best predicts predation risk depends on group movement

**DOI:** 10.1101/2021.05.25.445573

**Authors:** Poppy J. Lambert, James E. Herbert-Read, Christos C. Ioannou

## Abstract

Both empirical and theoretical studies show that an individual’s spatial position within a group can impact the risk of being targeted by predators. Spatial positions can be quantified in numerous ways, but there are no direct comparisons of different spatial measures in predicting the risk of being targeted by real predators. Here we assess these spatial measures in groups of stationary and moving virtual prey being attacked by three-spined sticklebacks (*Gasterosteus aculeatus*). In stationary groups, the limited domain of danger best predicted the likelihood of attack. In moving groups, the number of near neighbours was the best predictor but only over a limited range of distances within which other prey were counted. Otherwise, measures of proximity to the group’s edge outperformed measures of local crowding in moving groups. There was no evidence that predators preferentially attacked the front or back of the moving groups. Domains of danger without any limit, as originally used in the selfish herd model, were also a poor predictor of risk. These findings reveal that the collective properties of prey can influence how spatial position affects predation risk, via effects on predators’ targeting, hence selection may act differently on prey positioning behaviour depending on group movement.

## 1. Introduction

Animals often form groups to lessen their risk of predation [1, 2]. The risk of predation, however, is not distributed evenly across the different spatial positions an individual might occupy within the group. Risk can be higher for individuals on the group’s edge rather than in the centre (also known as marginal predation: [3-5]), for individuals positioned further from their near neighbours [6-8], or for those at the front or back of moving groups [9-11]. The majority of evidence for the different risk afforded by different spatial positions comes from observational studies of real predators targeting real prey, and from how prey respond by changing their position in groups in response to predatory attacks [7, 8, 12-14].

Measures to define prey position within groups fall into two broad categories: those describing centre-edge positioning, and those describing the degree of local crowding. Previous studies have tended to focus on one or the other, even though these measures tend to be correlated as individuals on the edge of groups have reduced local crowding. We are therefore limited in our understanding of which measures are particularly important in determining the predation risk faced by prey in different spatial positions. Such a comparison is challenging because it is difficult to accurately measure prey spatial positions during a sufficient number of, often unpredictable, attacks. Additionally, the spatial position within groups of real prey is known to be determined by a number of additional, confounding, factors, such as parasite load and boldness [15–19]. While models of virtual predators attacking simulated prey allow for unconfounded investigations of the effect of spatial position on risk [e.g. 20], these models have to make assumptions on how predators behave [6]. Finally, groups vary in their collective properties, such as whether groups are stationary or moving, and these collective properties can impact predation risk and foraging success [21-23]. However, whether such properties affect which measures of spatial position best predict predation risk is unknown.

To address these challenges, we presented simulations of virtual prey to three-spined sticklebacks (*Gasterosteus aculeatus*). Prey groups of 20 individuals were either stationary (with only small, uncoordinated ‘jitter’ motion) or had relatively slow directional movement (in addition to the jitter motion). The spatial positions of targeted and non-targeted prey were defined according to different measures previously used in the literature, and the success of each measure in predicting the likelihood of predation (i.e., attack) was assessed with an information criterion model comparison approach. By presenting real predators with virtual prey, the limitations of studies using predators attacking live prey, and simulated predators targeting simulated prey, can be overcome [3, 9, 24].

## 2. Material and methods

### (a) Subjects and housing

Three-spined sticklebacks (*G. aculeatus*) were caught from the River Cary in Somerton, UK (51.069990 latitude, −2.758014 longitude) in November 2017, and were transported by car to the University of Bristol. They were housed in 40 x 70 x 34cm (width x length x height) glass tanks on a flow-through recirculation system, with plastic shelters and plants for environmental enrichment, and kept at 14□ under a 11:13 light:dark cycle. Approximately 40 individuals were housed in each tank. The experiment was run from October – November 2018. The fish participating in the experiment were fed after each day of testing.

### (b) Experimental set-up

The testing arena (55 x 40 x 35 cm, L x W x H) was filled to a depth of 25 cm with water from the recirculating system that the fish were housed in. The walls of the back, side and bottom of the arena were covered with opaque white plastic. A screen made from a white translucent film (Rosco gel no. 252), was placed inside the front wall of the tank. When the simulated prey were projected onto this screen, they were visible to the predatory fish, while the predatory fish was also visible through the screen (figure 1). A projector (BenQ MW523) was positioned 82 cm in front and below the bottom level of the tank to minimise the bulb’s glare on the glass. Two strip lights were placed outside the tank behind the back wall to illuminate the arena from the rear. A camcorder (Panasonic VX870) was positioned directly in front of the tank, behind and above the projector. Videos were recorded at a resolution of 3840 × 2160 pixels at 25 frames per second. The entire experimental setup was enclosed by black curtains to visually isolate the experiment from the testing room.

**Figure 1:**
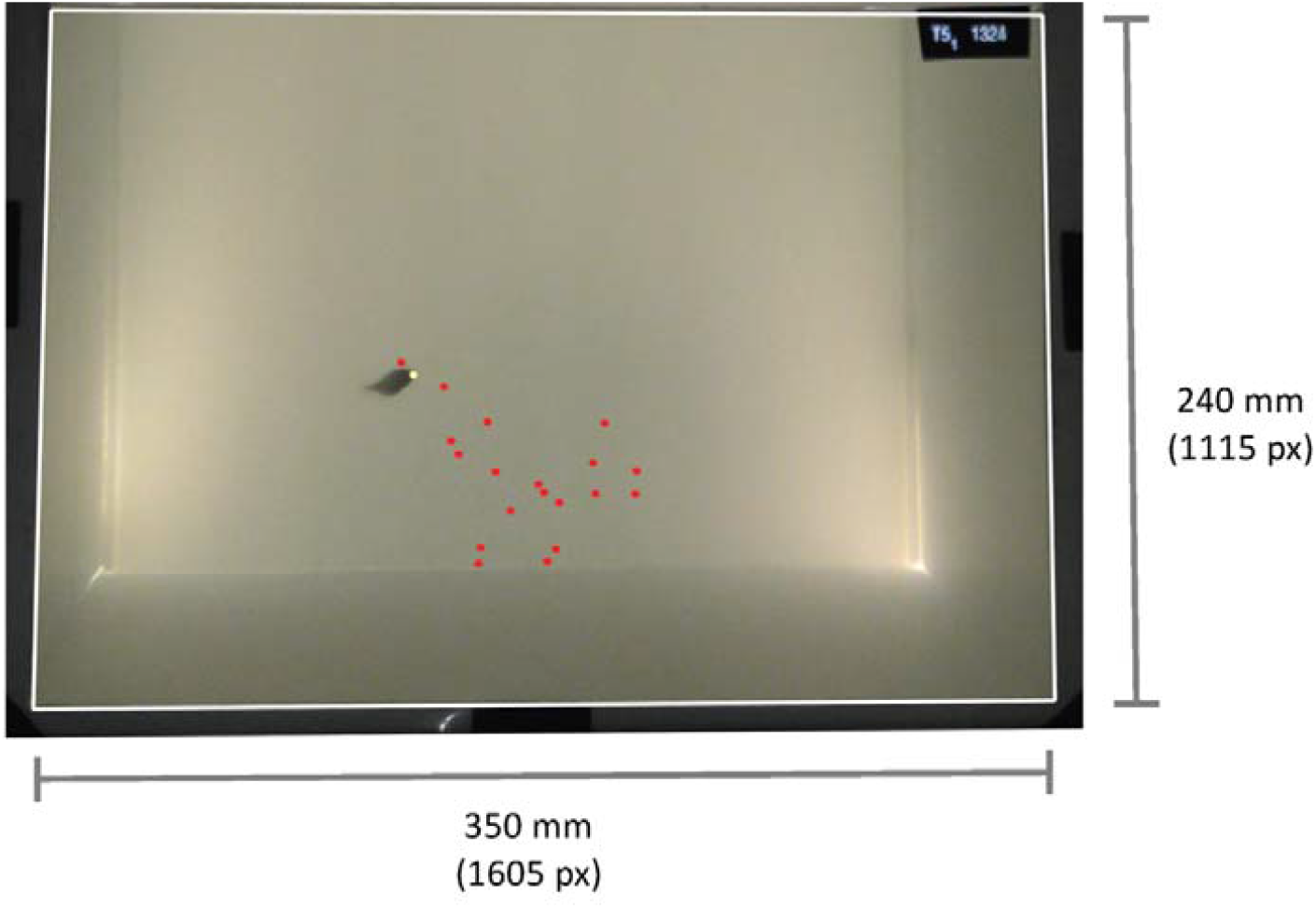
Experimental set-up. Frame from one of the experimental trials showing the moment a stickleback attacks one of the virtual prey. Virtual prey (red and yellow dots) have been overlaid on this image, which are not visible in the camera footage. The yellow prey indicates which prey was attacked, although all prey were projected as red dots in the trials. The white box represents the projection area where prey could appear. The black oblong in the top right-hand corner of the arena shows the trial number and the time stamp of the simulation. These time stamps were used to determine the locations of the prey in the calibration videos, where prey were visible.

### (c) Virtual prey simulations

We generated two types of prey simulations in MATLAB 2018b [25] (figure S1); stationary groups and moving groups (movies S1 and S2). Once projected onto the screen, the virtual prey appeared as red dots ∼ 3 mm in diameter. The prey could appear in an area of 350 x 240 mm, and in the moving simulations moved across the screen at 11 mm s^-1^. See electronic supplementary material for simulation details.

### (d) Prey presentation and identification

Each focal fish was presented with a unique prey simulation playing on a loop. Each loop lasted 40 seconds, with the first 10 s containing no prey, followed by 20 s of the prey being presented (the prey loomed in at the start over 1.67 s, and then loomed out over 1.67 s at the end of these 20 seconds), followed by 10 seconds without any prey, resulting in 20 seconds between each prey presentation. Each focal fish was exposed to the loop 15 times (∼ 10-minute trials), although we only used the first attack from each trial. A time stamp and playback identification number were shown in the top right corner of the simulations (but this was not visible to the fish as it was masked off from the fish’s field of view by black tape; figure 1). To infer which prey had been targeted in the trials, before each trial, we projected the identical simulation the fish was to receive but with prey projected as white dots. These ‘calibration’ videos allowed us to subsequently identify the locations of prey, so we could infer which prey had been targeted by matching the time stamps between the calibration video and the prey videos at the time of attack.

### (e) Experimental Procedure

Each subject participated in two trials: once in the stationary prey condition, and once in the moving prey condition. Fish were randomly assigned to one of two testing groups. One of the testing groups received the stationary prey condition first, and the other group received the moving prey condition first. The two trials for the same individual fish were separated by at least 24 hours. Within the moving prey condition, approximately half of the subjects were randomly assigned to receive the group moving from right to left (from the subject’s perspective) across the front of the tank, and the other half received prey moving left to right. Trials were generally alternated between the stationary and moving playbacks. Focal fish (n = 126) were given 10 minutes to acclimatise in the testing tank before any prey were projected.

### (f) Analysis and statistics

Only the first attack of each trial was used in the analysis [9]. An attack was defined when the stickleback orientated towards the screen and pecked (often more than once) at the screen. Where two prey were close to one another and we could not distinguish which of two (and in one trial, three) prey was the target, one prey was selected at random; this occurred in 8 of the 74 trials with an attack.

At the frame of the attack (i.e. when the fish made contact with the screen with its mouth open), we identified the time stamp of the simulation. We then manually tracked the position of all the prey in calibration videos at this time stamp using a bespoke script in MATLAB (2018b) [25]. This gave us the x and y coordinates (in pixels) of all the prey in the group. From these coordinates, we calculated the following measures of spatial position for all prey in each group: the Voronoi polygon area, limited domain of danger, nearest neighbour distance, number of near neighbours, distance to group centroid, whether or not the individual was on the convex hull, angle of vulnerability, and distance from the front of the group (the latter for moving groups only) (table 1). All were calculated using R version 3.6.0 [26]. The distribution of, and correlations between, these measures are shown in figure S2 (for stationary groups) and figure S3 (for moving groups).

**Table 1:**
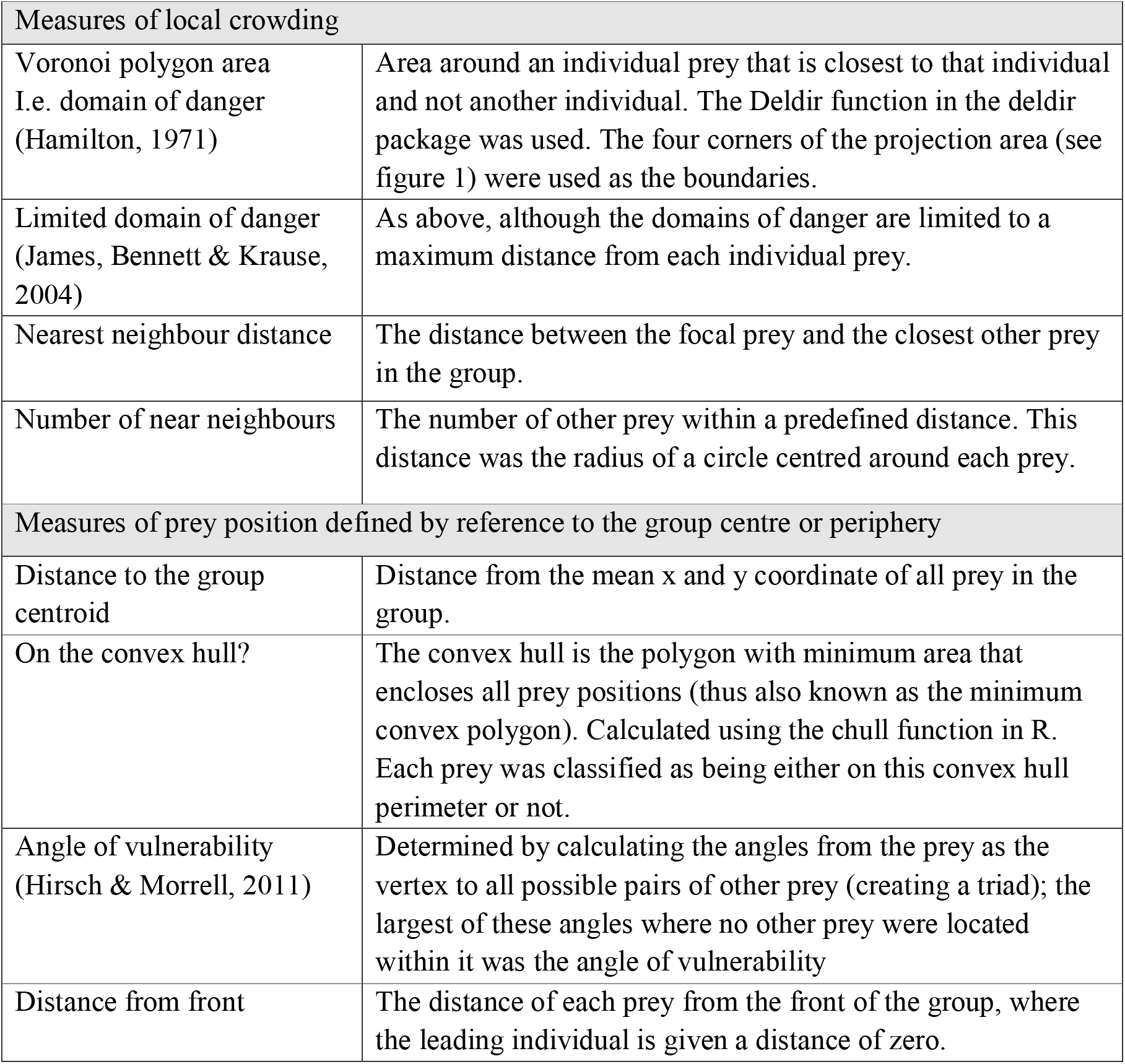
Description of the measures of spatial position used in our analysis. The distance from front measure was only applied to prey in moving groups.

To determine how well each measure of spatial position predicted the risk of an individual prey being targeted, we used binomial Generalised Linear Models (GLMs) with the default logit link function. The response variable was whether an individual was targeted (1) or not (0), and the explanatory variable was one of the measures of spatial position. A model for each measure of spatial position was constructed, separately for the stationary and moving groups; the data from the two conditions were analysed separately to avoid pseudoreplication, given that we could not keep track of fish identities across the two conditions. The likelihood of each model given the data was compared using the Akaike information criterion corrected for small samples sizes (AICc), as in [9]. A null model without an explanatory variable was also included to test whether the predictive power of models accounting for spatial position exceeded that of a null model after accounting for the extra parameter in these models.

The limited domain of danger and the number of near neighbours both require a maximum radius around the prey to be defined. For the limited domain of danger, this is the distance beyond which a predator will not attack the prey, even if that prey is the closest [27], and for the number of near neighbours, the distance within which near neighbours are counted [20]. We thus included in the AICc comparisons models for the limited domain of danger and number of near neighbours calculated using varying radius sizes, ranging from 10 to 500 pixels in 10-pixel increments. This parameter scan avoided any *a priori* or *post hoc* choice of the radius size, and allowed us to examine whether the radius size influenced the ability of the limited domain of danger and the number of near neighbours to predict predation risk.

## 3. Results

The test fish attacked the virtual prey in 35 trials with stationary prey (out of 123 trials) and in 39 trials with moving prey (out of 126 trials). In stationary groups, the targeted prey was best predicted by the limited domain of danger (LDOD) over a wide range of radius sizes (∼50 to 500 pixels); there was only a small range of radius sizes (∼50 to 100 pixels) where the number of near neighbours model outperformed it (specifically when LDOD radius sizes were smaller than 50 or greater than 300 pixels) (figure 2a). Greater local crowding was associated with a reduced predation risk (figure 3a). In contrast to the importance of the limited domain of danger in predicting risk, the Voronoi area (i.e. the domain of danger bounded by only the projection area) was a relatively poor predictor of being targeted (figure 2a, table S1). Measures of proximity to the group’s edge as predictors of risk were less well supported but were generally >2 AIC units less than the null model [28], suggesting that a prey’s angle of vulnerability, whether it is or is not on the convex hull edge, and the distance from the prey to the centroid still have explanatory power in predicting predation risk (figure 3a, table S1). For moving groups, the measures of spatial position that best predicted which prey were targeted were not consistent with the results from stationary groups. Although the number of near neighbours was the best predictor of risk when the radius size was optimised at 150 pixels, for much of the range of radius sizes examined, the angle of vulnerability and the distance to the group centroid (both measures of proximity to the group edge) had lower AICc values, i.e. were better predictors of risk (figure 2b, table S1). Whether prey were on the convex hull edge was less predictive of risk, but this had a lower AICc value than the remaining measures of local crowding (the limited domain of danger, nearest neighbour distance and Voronoi size (domain of danger)). Prey closer to the edge of the group were more likely to be targeted (figure 3b). The measure of spatial position relative to the front of the group (distance from the front), was a poor predictor of risk as it performed worse than the null model.

**Figure 2:**
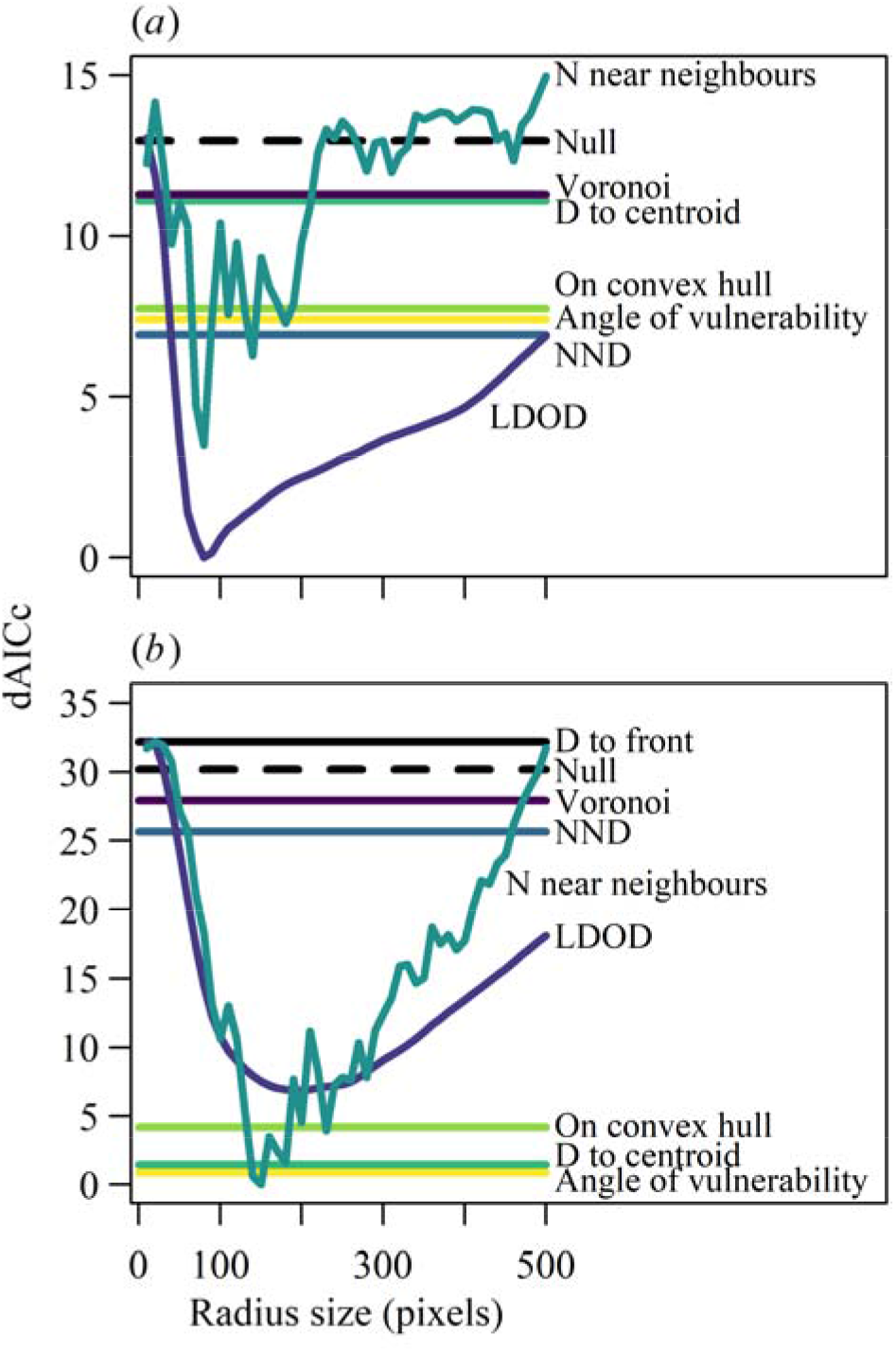
Model comparison results for stationary (a) and moving (b) groups based on the difference in the Akaike information criterion corrected for small sample sizes (dAICc) between the most likely model given the data (0 on the y axis) and each other model. In the model names, D represents distance, N to number, LDOD to the limited domain of danger and NND to nearest neighbour distance. As the dAICc for the limited domain of danger and number of near neighbours models are dependent on the radius size used to calculate these variables, dAICc values are plotted against radius size. For all other measures of spatial position, there is only one model (and hence dAICc value). Line colours for the different models correspond between the panels and figure 3.

**Figure 3:**
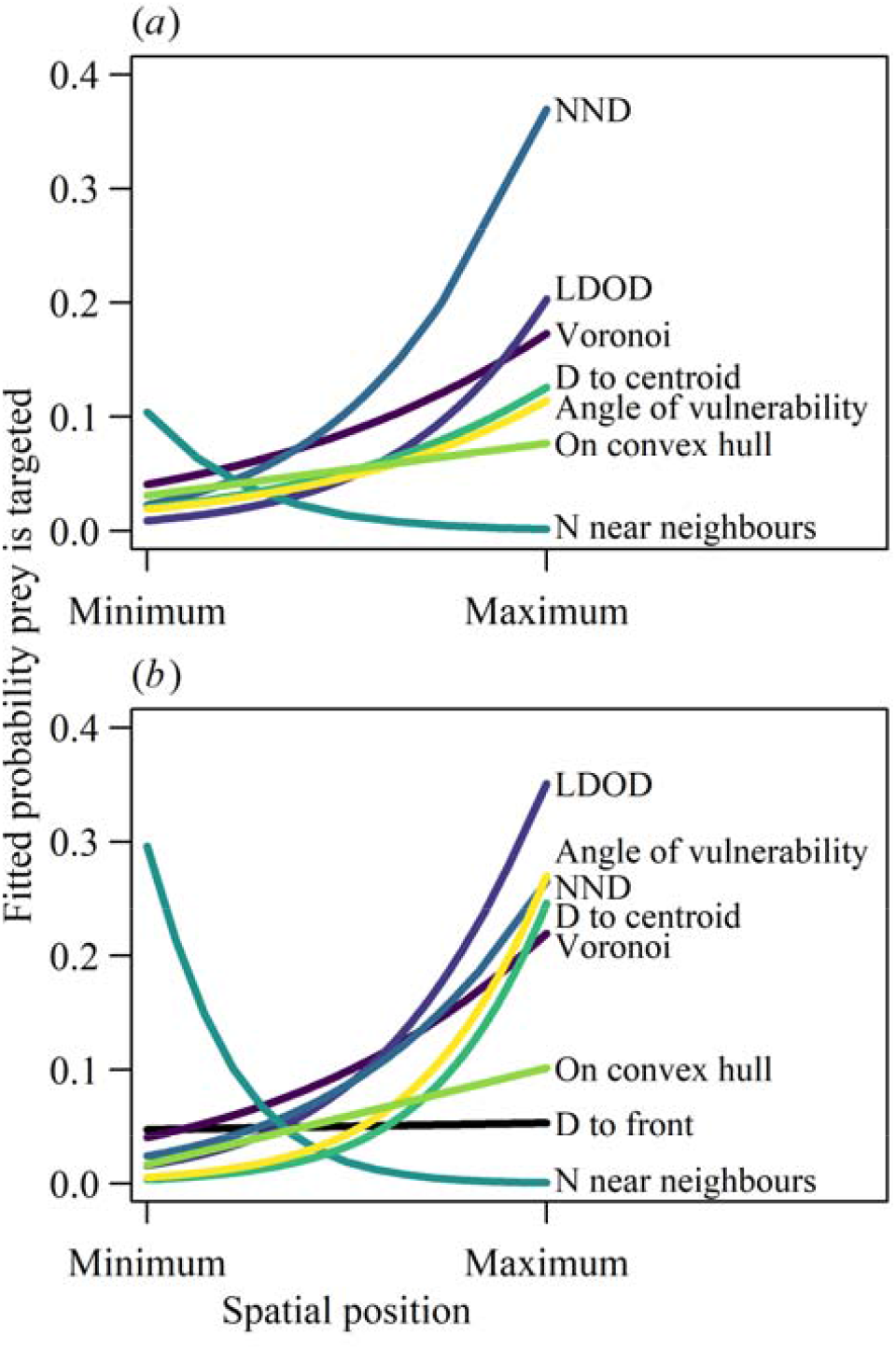
The fitted relationships between the probability that an individual prey in a stationary group (a) and moving group (b) is targeted and their spatial position, calculated using different measures, based on binomial GLMs. The radius size used for the LDOD and N near neighbours measures is the optimal determined by a parameter scan (see figure 2): 80 pixels for both measures in stationary groups and 200 and 150 pixels for LDOD and N near neighbours, respectively, in moving groups. As the scale of each measure of spatial position varies (shown in figures S2 and S3), the x axis ranges from the minimum to the maximum value of that measure observed in the data. The labels for the different measures of spatial position are arranged from top to bottom in order of the fitted probability of risk at the maximum value of the spatial measures. Colours of the lines of best fit correspond between the panels and figure 2.

## 4. Discussion

Our results demonstrate that even when comparing two relatively similar prey group types, where one group had directional movement and the other did not, targeting by predators was altered enough that the risk of being targeted was best predicted by different measures of spatial position. Measures of local crowding best predicted risk in stationary groups, with the limited domain of danger [27] outperforming all other measures. Although similar, the Voronoi area was a relatively poor predictor of risk, which lends substantial support to the more biologically-realistic limited domain of danger [27] in contrast to the unlimited domain of danger as originally formulated by Hamilton [6]. In moving groups, the best predictor of risk was the number of near neighbours but only for a small range of radius sizes within which neighbours are counted; more generally, measures of whether the prey were closer to the group edge performed best, rather than measures of local crowding. The radius that minimised the AICc for the limited domain of danger and number of near neighbours was substantially larger when groups were moving, possibly because moving objects have higher salience [29], which suggests that the radius size is context specific.

Animal groups vary in multiple aspects of their collective properties, such as the degree of directional movement, group shape, the spatial arrangement of individuals and the degree to which individuals change position within the group [30-33]. This can depend on species and ecological context [34-36], but can also vary over time within the same group and context [37, 23]. Given our results with groups which differed only with respect to their directional movement, multidimensional variation in collective properties of prey may make it difficult to generalise which measure of spatial position accurately predicts the risk of prey being targeted. This could help explain the surprising result in our study of no evidence that prey at the front of a moving group were more at risk, in contrast to other experiments with virtual [9] and real [10] prey. Further differences in the aspects of spatial position important in predicting risk may come from variation between predators, for example in the attack strategy of different species [38, 20] in hunger or experience between individuals of the same species [39], or from consistent personality differences between individuals within the same population [40].

During real-time observations of animal groups, counting the number of neighbours within a distance of a focal individual is a more versatile method of assessing spatial position than the Voronoi, limited domain of danger, distance to the centroid and angle of vulnerability measures, which rely on positional coordinates for all individuals in the group. However, like the limited domain of danger, the number of near neighbours requires the specification of the size of a radius around each prey. Without an *a priori* reason to set this radius at a particular size based on the biology of the predator and/or prey, a parameter scan is necessary, which would be difficult to do during real-time observations. Our parameter scans demonstrate that the likelihood of the models given the data (based on AICc values) for both the number of near neighbours and the limited domain of danger varied widely with radius size, demonstrating the importance of appropriately setting this distance. In contrast to the number of near neighbours for moving groups, however, the limited domain of danger outperformed all other measures of spatial position in stationary groups over a wide range of radius sizes, suggesting that the limited domain of danger is a robust and relatively accurate predictor of risk in stationary groups.

Our findings, where the predictive success of spatial measures differed for attacks on stationary groups and groups with directional movement, suggest that there may be no single best measure of spatial position for describing predation risk. Instead, the spatial measure that most predicts risk might be dependent upon the prey’s collective behaviour. This has implications for the selection pressure acting on prey in terms of the movement rules they should follow when locating themselves within the group [13, 41]. For example, our results suggest that when a group is relatively still, such as when they are resting or foraging within a patch, individuals should occupy areas which are more crowded to minimise their limited domain of danger, regardless of whether this area falls near the edge of the group. However, occupying the optimal spatial position within the group may be constrained by cognitive abilities in determining where to move to, and the behaviour of other individuals in the group [42-44].

## Acknowledgements

This work was supported by Natural Environment Research Council Independent Research Fellowship NE/K009370/1 and Leverhulme Trust Grant RPG-2017-041 V awarded to C.C.I.. J.H-R. was supported by the Whitten Lectureship in Marine Biology and a Swedish Research Council grant: 2018-04076.

