## Supplementary material for "The measure of spatial position within groups that best predicts predation risk depends on group movement"

1 Figure S1 and Simulation Details
2 Figure S2 and S3
3 Table S1
4 Supplementary Dataset Information
5 Supplemental Material References

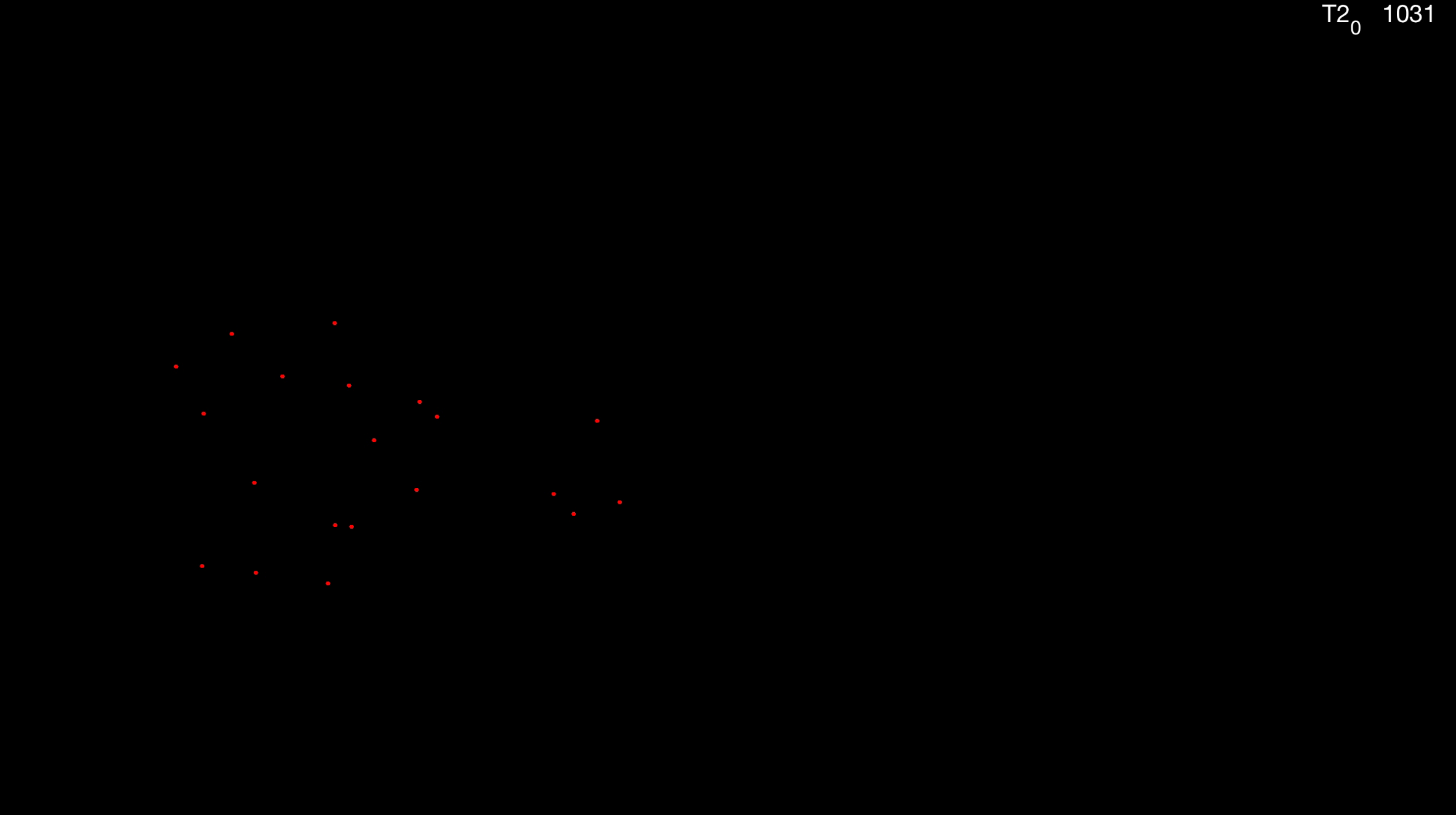

Figure S1: 20 virtual prey (red dots) simulated in MATLAB 2018b.

Simulation Details

For both simulation types, the relative positions of each of the 20 prey were randomly selected from within a circular radius of 75 pixels. For the stationary prey, the initial starting location (x and y location of where the prey would appear on the screen) was randomly chosen within the boundaries of the projection and added to the relative spatial positions. For the moving groups, the initial y location of the prey was again randomly chosen (and added the prey’s relative y coordinates), but for the x component, we added a linear component to the prey’s relative x coordinates so that either appeared on the left of the arena and moved to the right, or appeared on the right of the arena and moved to the left. When projected, this made the prey move across the screen at 11 mm s^-1^. We also added small jitter motion to the prey of both stationary and moving groups, making them appear more lifelike. To do this, prey moved on a correlated random walk (0.05 pixels per timestep) choosing a direction of motion from within ± 30 degrees of their previous direction. If prey moved more than five pixels from their initial starting point, they turned towards their initial starting point, ensuring that for the stationary groups, each prey remained within five pixels of their initial position, and for the moving groups, prey remained within five pixels of their initial y coordinate. Because groups in nature often appear oblong in shape (i.e. longer in one dimension than another) (Hemelrijk & Hildenbrandt, 2008), we applied an aspect ratio to the simulations so that the width of the group was on average 1.5 ± 0.2 greater than the height of the group (mean ± 1 SD).

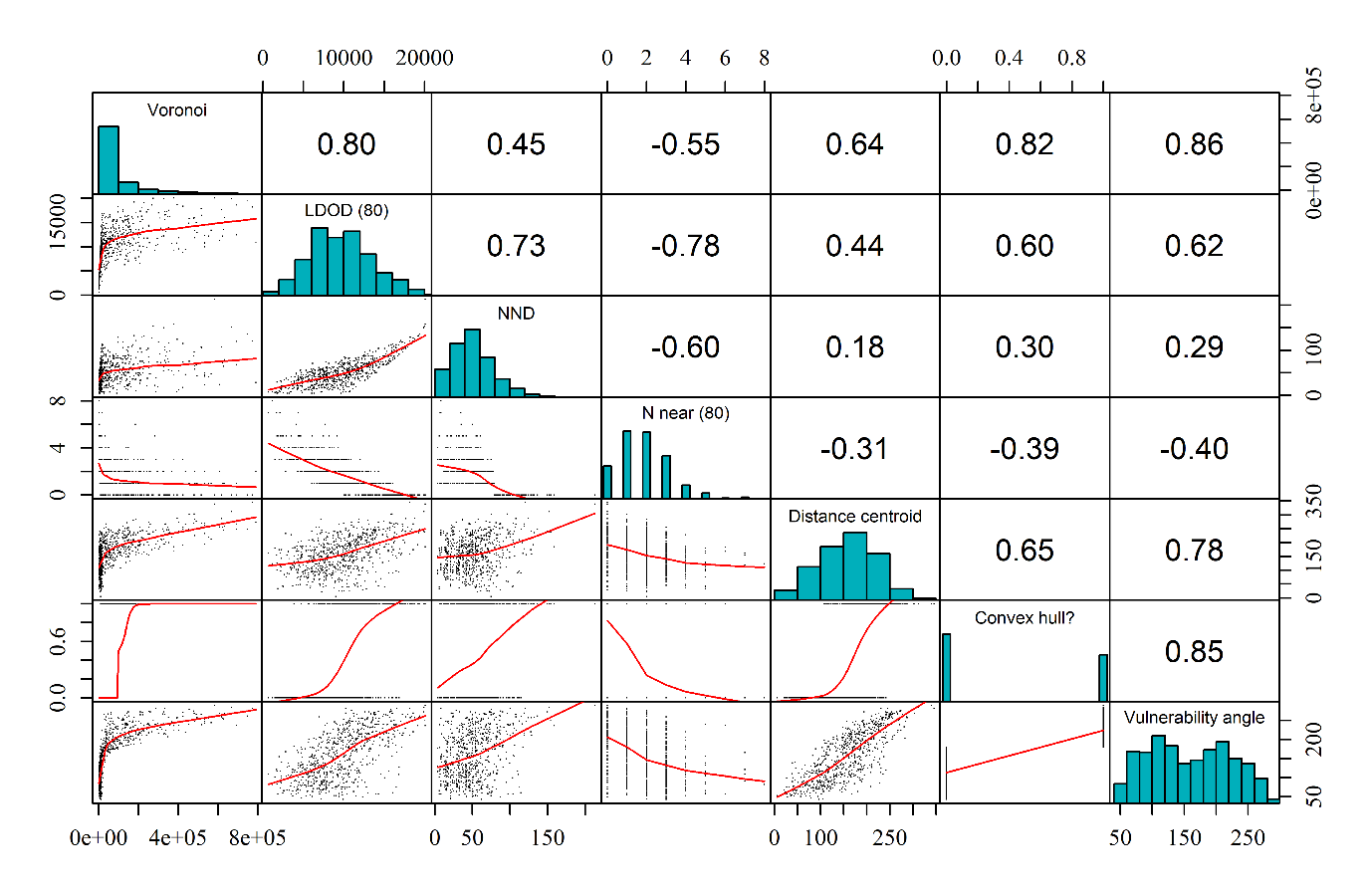

Figure S2: The distributions of, and correlations between, the seven measures of spatial position in the stationary virtual prey groups. Voronoi is the area of the domain of danger, LDOD is the area of the limited domain of danger, NND is nearest neighbour distance, N near is number of near neighbours, Distance centroid is the distance to the group centroid, Convex hull? is whether the individual was one of the vertices of the convex hull around the group, and Vulnerability angle is the size in degrees of the angle of vulnerability. The value in brackets after LDOD and N near denote the radius size in pixels used to calculate those measures of spatial position. Distances are given in pixels and areas in pixels squared. Correlation coefficients (top right cells showing single values) are Spearman’s r_s_. The red curve in the scatterplots are LOWESS smoothed curves.

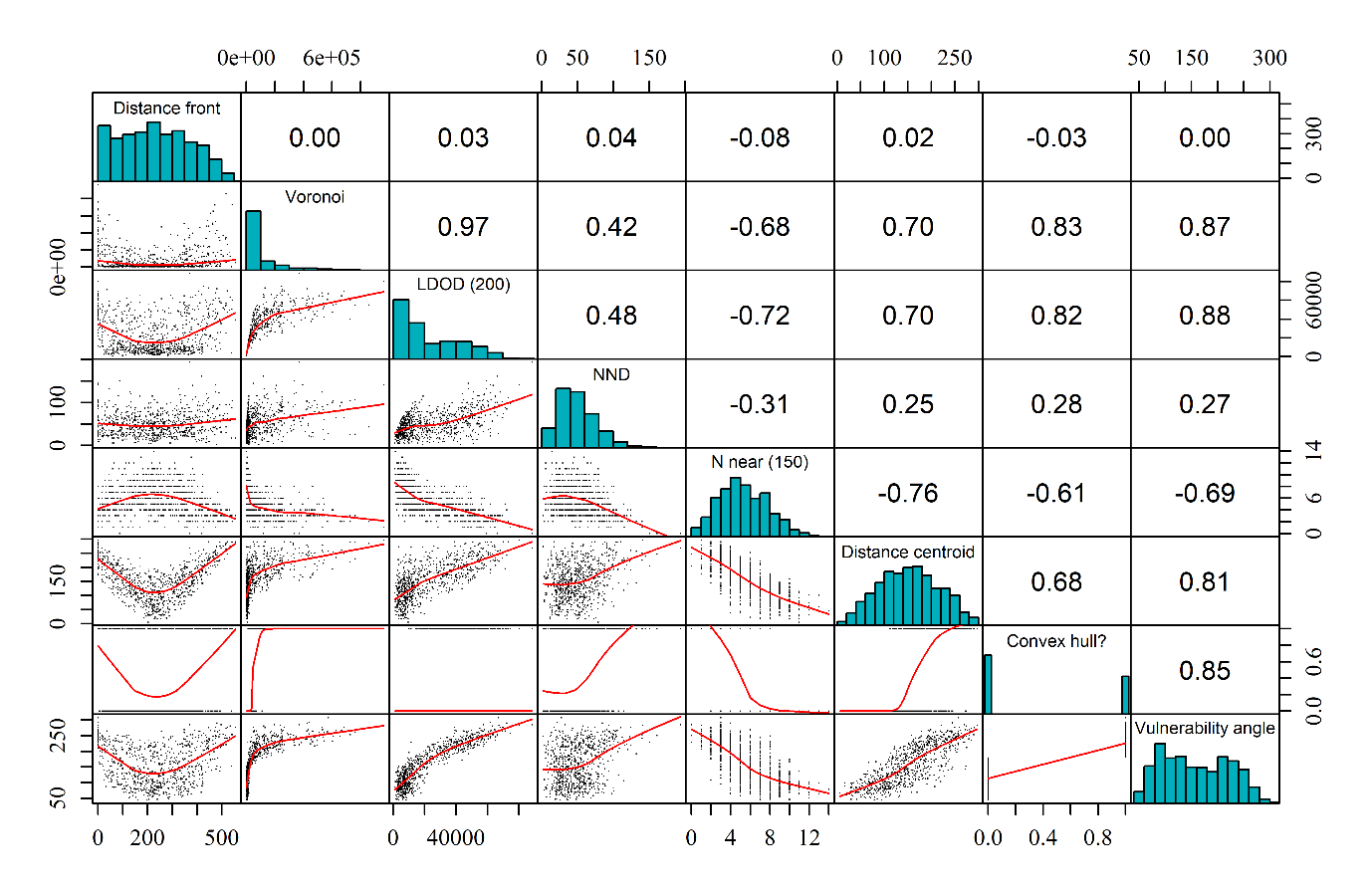

Figure S3: The distributions of, and correlations between, the eight measures of spatial position in the moving virtual prey groups. Distance front is the distance from the individual to the front of the group, Voronoi is the area of the domain of danger, LDOD is the area of the limited domain of danger, NND is nearest neighbour distance, N near is number of near neighbours, Distance centroid is the distance to the group centroid, Convex hull? is whether the individual was one of the vertices of the convex hull around the group, and Vulnerability angle is the size in degrees of the angle of vulnerability. Distances are given in pixels and areas in pixels squared. The value in brackets after LDOD and N near denote the radius size in pixels used to calculate those measures of spatial position. Correlation coefficients (top right cells showing single values) are Spearman’s r_s_. The red curve in the scatterplots are LOWESS smoothed curves.

Table S1: Model comparison results based on the difference in the Akaike information criterion corrected for small sample sizes between the most likely model given the data and each other model (dAICc). d.f. is the degrees of freedom. Models are ordered from the most to least likely. For brevity, only the best performing (lowest dAICc) models for the limited domain of danger and number of near neighbours are included, with the radius size (in pixels) that optimises their predictive power given in brackets. A radius size of 80 pixels maximised the power (i.e. minimised the AICc) of the limited domain of danger and the number of near neighbours to predict risk in stationary groups, while larger radii of 200 pixels and 150 pixels maximised the predictive power of the limited domain of danger and number of near neighbours, respectively, in moving groups.

| Explanatory variable | dAICc | d.f. |
| --- | --- | --- |
| *Stationary groups* |  |  |
| Limited domain of danger (80) | 0 | 2 |
| Number of near neighbours (80) | 3.5 | 2 |
| Nearest neighbour distance | 6.9 | 2 |
| Angle of vulnerability | 7.4 | 2 |
| Is prey on convex hull? | 7.7 | 2 |
| Distance to centroid | 11.1 | 2 |
| Voronoi size (domain of danger) | 11.3 | 2 |
| Null model | 13 | 1 |
| *Moving groups* |  |  |
| Number of near neighbours (150) | 0 | 2 |
| Angle of vulnerability | 0.9 | 2 |
| Distance to centroid | 1.5 | 2 |
| Is prey on convex hull? | 4.2 | 2 |
| Limited domain of danger (200) | 6.9 | 2 |
| Nearest neighbour distance | 25.7 | 2 |
| Voronoi size (domain of danger) | 27.9 | 2 |
| Null model | 30.2 | 1 |
| Distance from the front of the group | 32.2 | 2 |

Supplementary Dataset Information

Data from the trials where the fish made at least one attack. The x and y coordinates for each prey at the moment of the first attack per trial are given, the attacked prey individual is specified, as is the coordinates of the corners of the projected area. From these data, the measures of spatial position are calculated for each prey.

MATLAB (R2018b). Natick, Massachusetts: The MathWorks Inc.; 2018.
